# On the clinical benefit of on-scalp MEG: A modeling study of on-scalp MEG epileptic activity source estimation ability

**DOI:** 10.1101/2022.01.28.478237

**Authors:** Karin Westin, Sándor Beniczky, Matti Hämäläinen, Daniel Lundqvist

**Affiliations:** NatMEG, Department of Clinical Neuroscience, Karolinska Institutet, Stockholm, Sweden; Clinical Neurophysiology, Karolinska University Hospital, Stockholm, Sweden; Department of Clinical Neurophysiology, Aarhus University Hospital, Denmark and Danish Epilepsy Centre, Dianalund, Denmark; Department of Psychiatry and the Athinoula A. Martinos Center for Biomedical Imaging, Massachusetts General Hospital, Harvard Medical School, Charlestown, MA, 02129, USA

**Keywords:** Epilepsy, clinical, on scalp magnetoencephalography, modeling

## Abstract

**Objective:** Whole-head on scalp magnetoencephalography (osMEG) is a novel, cutting-edge functional neuroimaging technique that positions MEG sensors closer to the cortical sources. OsMEG allows both for free head movements and improved spatial resolution compared to conventional MEG. OsMEG thus might improve clinical epilepsy evaluations. However, it remains largely unknown how osMEG characterizes epileptic activity. Here, we aimed to compare epileptic activity source estimation accuracy of osMEG, high-density EEG (hd-EEG), conventional MEG (convMEG) and subdural EEG (sbdEEG).

**Method:** IED and seizure onset zone source estimations of osMEG, hdEEG, convMEG and sbdEEG were evaluated using equivalent current dipoles. Cancellation index of all non-invasive modalities were calculated and compared statistically. To further investigate any similarity between osMEG and sbdEEG, representational similarity analysis was used to compare IED source estimations of these two modalities.

**Results:** We found that osMEG IED source estimations were significantly (p<0.05) better than both convMEG and hdEEG. Furthermore, osMEG mesial temporal lobe SOZ source estimations were superior to those of convMEG and hdEEG. OsMEG cancellation index did not differ significantly from convMEG. Interestingly, comparing osMEG and sbdEEG IED source estimation demonstrated that osMEG might be less sensitive to source directions than convMEG.

**Conclusion:** We demonstrated that whole-head osMEG exhibited very accurate non-invasive IED and SOZ source estimations, better than both hd-EEG and convMEG.

**Significance:** OsMEG has a promising potential to become a safe, highly sensitive neuroimaging modality for whole head epilepsy evaluations.

**Highlights:**

- Analysis of novel on-scalp MEG (osMEG) sensors in epilepsy evaluations
- Compares osMEG, EEG, conventional MEG & intracranial EEG epilepsy source estimations
- Our study demonstrates potential great clinical value of osMEG whole-head sensors

## 1. Introduction

On scalp magnetoencephalography (osMEG) is a novel variant of the non-invasive MEG technology that enables MEG sensors to be placed directly on the scalp, instead of in a helmet, as is done in conventional MEG (convMEG) (Boto et al., 2018; Xie et al., 2017). OsMEG sensor technology today includes both highTc SQUIDS (Zhang et al., 1993) and optically pumped magnetometers (OPMs) (Budker and Romalis, 2007). Although both OPMs and highTc SQUID sensors offer similar spatial resolution and sensor-to-scalp distance, whole head sensor coverage can currently be achieved only using OPMs (Boto et al., 2021), which are also as of now commercially available from several manufacturers (QuSpin, FieldLine, Kernel, Cerca Magnetics). Since this study focuses on *whole-head osMEG* measurements, we therefore focus on OPMs.

With OPMs, the positioning of sensors can be adapted to the individual head shape, giving a sensor-to-scalp distance of 5 – 7 mm. This is significantly smaller than the typical distance of in convMEG which offers a 20-40 mm sensor-to-scalp distance (Heiden, 1991; Iivanainen et al., 2017; Riaz et al., 2017). Furthermore, in convMEG the head must remain within the helmet-shaped sensor array and any head movements have to be compensated for using computational techniques (Taulu and Simola, 2006). With osMEG, sensors can be attached to a wearable helmet, thus allowing movements during measurements without significant reduction of signal-to-noise ratio and without the need of computational movement compensation (Boto et al., 2018, 2021).

Today, convMEG is routinely used for localization of interictal epileptiform discharges (IEDs) (De Tiège et al., 2017; Hari et al., 2018). Notwithstanding, ictal discharges have also been recorded with convMEG. In practice however, EEG is currently the only method routinely used for long term monitoring and seizure registrations (Jayakar et al., 2008, 2014) because EEG arrays allow free movement during longer measurements. OsMEG has the potential of becoming the first MEG technology which allows for long-term monitoring, and hence ictal MEG measurements. This development has the potential to lead to important progress in epilepsy evaluations. More importantly, estimation of the seizure onset zone from scalp EEG can be inaccurate because of limits in EEG sensitivity and accuracy, requiring complementary intracranial recordings (Rossi Sebastiano et al., 2020). While optimally placed intracranial EEG electrodes can pinpoint the sources of epileptic activity accurately, only limited number of subregions of the brain can be mapped. In addition, the invasive technique bears additional risks for adverse events compared to the non-invasive methods (Chassoux et al., 2018). Due to the enhanced sensitivity and spatial precision gained by osMEG’s improved sensor-to-brain proximity relative to convMEG (Schneiderman, 2014; Xie et al., 2015; Boto et al., 2016; Iivanainen et al., 2017) an osMEG sensor array can enable recordings of whole-head seizure data of significantly higher quality than what is possible today.

Several simulation studies have indicated that the reduced sensor-cortex distance drastically increases both information content and spatial resolution of the osMEG signals, as compared to convMEG (Schneiderman, 2014; Boto et al., 2016; Iivanainen et al., 2017; Riaz et al., 2017). In agreement with these theoretical predictions, experimental studies have confirmed that osMEG sensor systems increase signal amplitude, spatial resolution and accuracy (Andersen et al.; Xie et al., 2017; Boto et al., 2021). Importantly, in the first osMEG measurement on an epilepsy patient, our group has recently demonstrated that osMEG is indeed able to detect low amplitude spikes which were missed by both convMEG and scalp EEG: we identified twice as many IEDs as convMEG and three times as many IEDs as EEG (Westin et al., 2020). Several lines of evidence thus indicate that osMEG has a strong potential to become an excellent tool for non-invasive seizure onset localization, beyond the current state-of-the-art of conventional neurophysiological techniques.

However, there are still many unknowns regarding how well osMEG can detect and locate epileptic activities. Here, we present a modeling study that evaluates the capability of osMEG to detect as well as estimate the source of simulated seizures and IEDs. We compare *osMEG* to *convMEG*, to high-density EEG (*hdEEG*), and to subdural EEG (*sbdEEG*). Our hypothesis is that sbdEEG has the best capability to detect and estimate epileptic activity, followed in order by osMEG, convMEG and hdEEG. Potentially, the difference between sbdMEG and osMEG is quite small, and our aim is hence to help estimate the ultimate potential benefits provided by this novel recording technique by modelling the degree of similarities and dissimilarities between osMEG, sbdEEG, convMEG and hdEEG, and to provide valuable design input for osMEG device and protocol development.

For this work we use realistically modeled epileptic activity and have divided the study into four different sections:

1. *Localization of the irritative zone*. Routine epilepsy evaluations utilize non-invasive convMEG and hdEEG to estimate IED sources. To investigate the clinical potentials of osMEG, we compared IED source estimations of all three non-invasive modalities including a whole-head sbdEEG grid.
2. *Localization of the epileptogenic network*. Clinical presurgical evaluations aim to estimate the seizure onset zone (SOZ) of gradually propagating epileptic activity (Bartolomei et al., 2017). Non-invasive techniques are often used to delineate potentially seizure generating brain regions to be subsequently mapped by stereo-EEG. In order to characterize a potential role for osMEG within presurgical planning, we performed non-invasive source estimations of spatio-temporally evolving seizure activities. In addition, osMEG SOZ source imaging was compared to the model of a whole-head intracranial EEG montage.
3. *Cancellation index*: Since all epileptic foci are spatially extended and the cortical surface is folded, there are different source orientations within the cortical epileptic focus. The contributions of sources of different polarities might thus cancel out, which is detrimental for the characterization of the extent of the epileptic foci. In order to investigate epileptic source cancellations in osMEG, we compared its cancellation index (Ahlfors et al., 2010) with hdEEG and convMEG.
4. *Similarity of osMEG and sbdEEG*. To further analyze the potential osMEG, we compared the IED source estimation ability of whole-head osMEG array to an intracranial sbdEEG whole-head grid.

## 2. Materials and methods

Two types of epileptic activity (IEDs, and seizure activity) were simulated. The resulting sensor data from osMEG, convMEG, hdEEG and sbdEEG, were modeled, followed by analysis of epileptic activity and source estimation.

### The anatomical model

The anatomical model in the sample dataset of the MNE Python software package was used (Gramfort et al., 2013). The model includes an original T1 MPRAGE sequence MRI recorded at a 1.5 Tesla Siemens scanners. A full segmentation of the head and brain was performed using Freesurfer (Dale et al., 1999; Fischl et al., 1999). Skin, skull, and brain surface boundaries were created using MNE Python (Gramfort et al., 2013).

### 2.1 The forward models

#### Source space

The potential sources were constrained to the cortical mantle using the cortical surface reconstruction provided by Freesurfer. The source orientation was constrained to be normal to the cortex.

#### OsMEG, ConvMEG

The gain matrix for all sources was computed using a single-compartment (intracranial volume) boundary-element model (BEM) in MNE Python. The sensor definitions were provided by the coil_def.dat file in MNE Python.

#### HdEEG

A three-compartment BEM (skin, skull, and intracranial volumes) was used in the computation of the gain matrix in MNE Python.

#### SbdEEG

Brain surface boundaries as well as a one-layer compartment volume conductor model taking only the inner skull layer into account, a source space and an EEG forward model was computed using MNE Python. A sbdEEG forward model was computed using MNE Python EEG forward solution with a modified one-layer (only inner skull) BEM model. (Gramfort et al., 2013).

### 2.2 Sensor arrays

#### OsMEG

The sensors were modeled as optically pumped magnetometer (OPM) sensor cubes with side length 5 mm. The sensor output was computed by integration over the sensing volume. Noise levels were set to 6 fT/Hz^(1/2)^ in accordance with other osMEG simulation studies (Iivanainen et al., 2017; Boto et al., 2018) utilizing optimal, best-case noise levels. An array of 128 sensors were modeled and placed with 1 cm intersensor distance over the scalp with 1 mm standoff, also in accordance with other osMEG simulation studies (Iivanainen et al., 2017).

#### ConvMEG

We employed the sensor array configuration of the 306-channel Vectorview system (MEGIN OY, Helsinki, Finland). The sensor output was determined by integration over the pickup coil plane and noise levels were set to 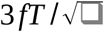 for both the planar gradiometers and magnetometers of the system, respectively.

#### HdEEG

We employed a 128-electrode montage and evaluated the electric potential at the center of each electrode. The montage was based upon a standard 128-electrode EEG cap (REF)

#### SbdEEG

Standard human subdural EEG electrode center-to-center distance of 5 mm was used, resulting in 223 electrodes across each hemisphere.

### 2.3 Simulation of epileptic and background activities

Epileptic activities include seizure (also called ictal) activity and IEDs. Seizures are often characterized by an initial low-voltage, high-frequency LVHF-activity) activity followed by sustained, repetitive sharp transient activity (sustained seizure activity) (Tao et al., 2007b; Jenssen et al., 2011). IEDs consist of transient, sharp, often isolated activities in between seizures. HFLA activity was modeled as a 25 Hz sine function. Both sustained seizure activity and IEDs were modeled using the Wendling model (Wendling et al., 2000; Wendling et al., 2002). This model is a further development of the Jansen and Rit neural mass model (Jansen and Rit, 1995). Here, a set of coupled differential equations are used to model a cortical column of pyramidal cells receiving both inhibitory and excitatory feedback. The output, the summed post-synaptic potential of the neural population, can be tuned to simulate both alpha activity and evoked potentials depending on parameter setting. Within the Wendling model, parameters reflecting synaptic coupling and excitation/inhibition balance have been modified to further resemble epileptic cortical tissue characteristics, resulting in an epileptic activity-generating neural mass model. The model has been validated in several studies. The Wendling model was used to simulate both sustained seizure activity, and interictal epileptic spikes (see Fig 1 for an example of simulated IEDs) (Wendling et al., 2000, 2002, 2005; Cosandier-Rimélé et al., 2010).

**Figure 1:**
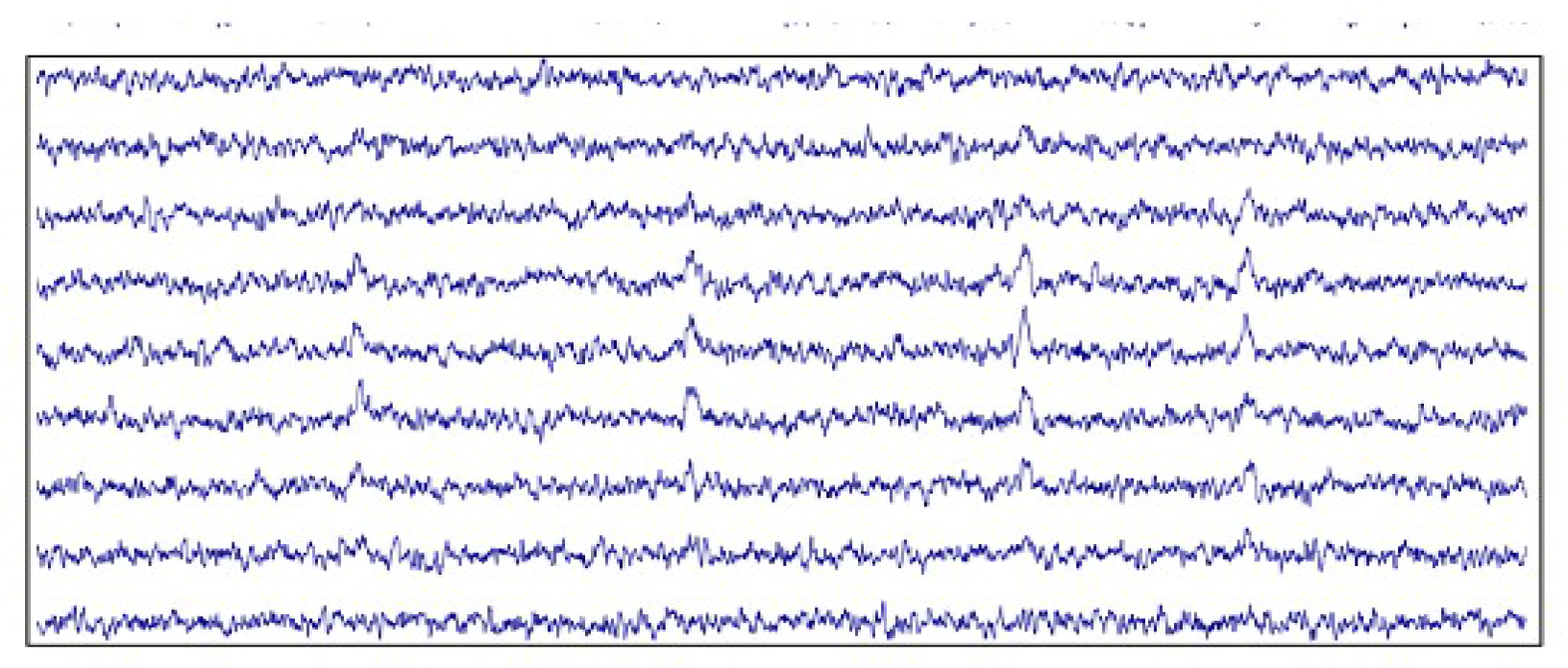
Simulated interictal epileptiform discharges (IEDs) in on scalp MEG (osMEG) sensor data. Example of IEDs modeled using the Wendling model simulated as raw osMEG sensor data.

#### Background activity

Routine clinical neurophysiological epilepsy evaluations require the epileptologist to discriminate both IEDs and seizure activity from the background brain activity. Thus, in order to simulate realistic epileptic evaluations, brain noise was added to all channels of the non-invasive modalities. In hdEEG, noise levels were set to achieve a baseline amplitude of 20-40 microV to resemble average adult alpha activity. Correspondingly, MEG brain noise levels were set to achieve baseline amplitude 200 fT (Kane et al., 2017; Rampp et al., 2020).

### 2.4 Source estimation

Clinical neurophysiological epilepsy evaluations aim at detecting either epileptic foci (irritative zone) generating IEDs, or the seizure onset zone and propagation (epileptogenic network) by localization of LVHF -activity. We modeled both of these situations in *2.4.1 irritative zone localization* and *2.4.2 seizure network localization*.

#### 2.4.1 Irritative zone localization

Eight sites (placed in mesial temporal lobe, lateral temporal lobe, frontopolar region, lateral frontal lobe, central region, parietal lobe and occipital lobe) were selected as epileptic foci (see Figure 2 for placement of these sites). Gradually growing foci were created by increasing the radius by 1 millimeter (mm) from radius 1.5 mm (area = 0.07 cm^2^) to radius 60 mm (area = 113 cm^2^). Each foci were set to generate interictal epileptic spikes using the Wendling model described in (see *2.3 Simulation of epileptic and background activities*). Dipole densities generated in physiological, non-epileptic settings lie within the range *q* = 0.16-0.77 nAm/mm^2^ (Murakami and Okada, 2015). The corresponding values of spike-producing cortical tissue are unknown. The intracellular event, the paroxysmal depolarizing shift, underlying interictal epileptic spikes results in a large long-lasting membrane depolarization (Ayala et al., 1970) and might result in a larger current dipole moment densities than normal pyramidal cell post-synaptic potentials. To accommodate for this possibility, dipole strength of the simulated interictal epileptic spikes were set to *q* = 0.77 nAm/mm2. For each focus, a total of 20 isolated interictal epileptic spikes were generated together with background noise activity as described in *2.3 Simulation of epileptic and background activities*. Raw data were simulated for osMEG, convMEG, sbdEEG and hdEEG. Clinical IED source localization is based upon visual inspection of raw data. In order to resemble the clinical setting, an experienced clinician inspected the sensor data to characterize IED visibility in all modalities. The equivalent current dipoles (ECDs) (Sarvas, 1987) were hereafter estimated from both single interictal epileptic spikes events, as well as averaged interictal epileptic spikes. The Euclidean distance between the origin as well as the perimeter of the foci and the ECD was determined for all foci sizes.

**Figure 2:**
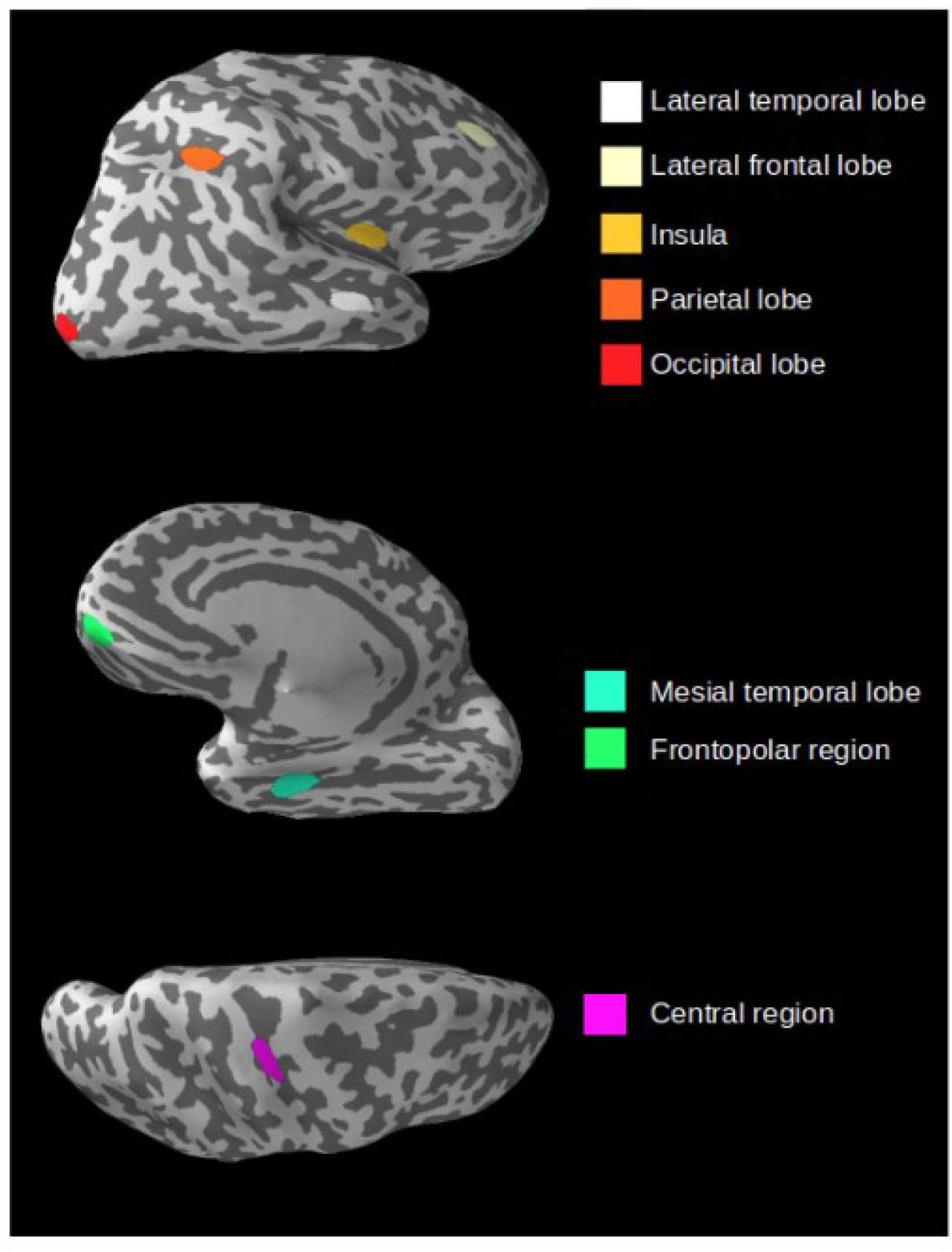
Placement of irritative zones. Localization of the eight anatomical sites selected as epileptic foci. Note that the zone site 3 cm^2^ utilized here was only to visualize the sites.

#### 2.4.2 Seizure network

In order to resemble common seizure types, one temporal lobe seizure and one frontal lobe seizure was simulated. The temporal lobe seizure propagated to ipsilateral prefrontal cortex, contralateral prefrontal cortex and contralateral temporal lobe. As studies have demonstrated contralateral seizure propagation times between 1-47 seconds, propagation time to contralateral hemisphere was set to 3 seconds (Jenssen et al., 2011). The frontal lobe seizure originated from right hemisphere lateral frontal lobe and propagated to homotypic regions on the contralateral lateral frontal lobe within four seconds (Blume et al., 2001). Each seizure was initiated by three seconds of LVHF -activity succeeded by sustained epileptic activity. Initial seizure onset size was set to area with radius 8 mm (area 2 cm^2^) (Ray et al., 2007; Tao et al., 2007a). The seizures grew gradually from radius = 6, 8 mm, 12 mm, 15, 20, 25, 30, 35 and, 40 mm. Hereafter, a forward model for sbdEEG, osMEG, convMEG and hdEEG was computed and sensor data simulated as described above. To resemble a typical clinical setting, the sensor data of each modality was inspected by an experienced clinician. The first visually detectable seizure activity was identified for each modality. The ECD was determined at the time of the first detectable seizure activity. For both simulated seizures, the Euclidean distance between the fitted dipoles for each modality and the simulated seizure origin was calculated.

### 2.5 Cancellation Index

Cancellation index aims to characterize the loss of non-invasive EEG/MEG signals due to simultaneous activation of sources with opposing polarity. Cancellation index was only computed for hdEEG, convMEG and osMEG. In this study, we utilized the metric defined in Ahlfors 2010 (Ahlfors et al., 2010):

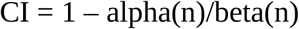

where alpha(n) = [Sigma_j_^M^(Sigma_j_^n^a_ij)^2^]^1/2^ and

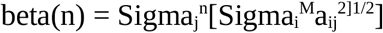

Gradually growing epileptic foci were determined as in *Source estimation*. Average cancellation index for all modalities were determined for all foci. Student’s T-test was used to determine any statistically significant difference between both foci location, and modalities.

### 2.6 Comparison of osMEG and sbdEEG

In order to compare osMEG and sbdEEG source estimation ability of epileptic activity depending on source depth and orientation, we employed an osMEG - sbdEEG similarity metric (*dissimilarity score*) based upon a modified representational similarity analysis (Kriegeskorte et al., 2008).

OsMEG and sbdEEG forward modeling was conducted as described in *The forward models*. Hereafter, the right hemisphere cortex was subdivided into 4096 patches with radius 5 mm. At each patch, ten IEDs with dipole strength 10 nAm were simulated and epochs of length 0.2 seconds were created centered around the IED peak. An inverse solution of the averaged IED epochs in osMEG sensor data was calculated using MNE Python dSPM with signal-to-noise ratio = 3 and regularization parameter = 1/9 (Gramfort et al., 2013). No inverse solution of the averaged sbdEEG IED epochs were computed as routine clinical sbdEEG analyses are based upon visual inspection of raw sensor data only. The *dissimilarity score* was now computed for each patch according to the following:

1. Two [223×1]-arrays containing sbdEEG (*sbdArray*) and osMEG (*osArray*) data were created according to the following:
  *sbdArray*: Contains averaged IED epoch sbdEEG sensor data from all 223 right hemisphere sbdEEG electrodes.
  *osArray*: The osMEG dSPM source estimate was extracted from 223 virtual sensors positioned at the right hemisphere sbdEEG electrodes.
2. For each array, a [223×223]-dissimilarity matrix containing the pairwise Euclidean distance between all array entries was created:

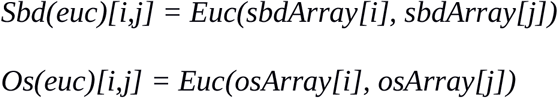
3. To determine the source estimation capability of each modality, the *peak activity channel* was defined as the channel with the longest Euclidean distance to all other channels. Hereafter, the Euclidean distance between the source and the *peak activity channels* was determined:

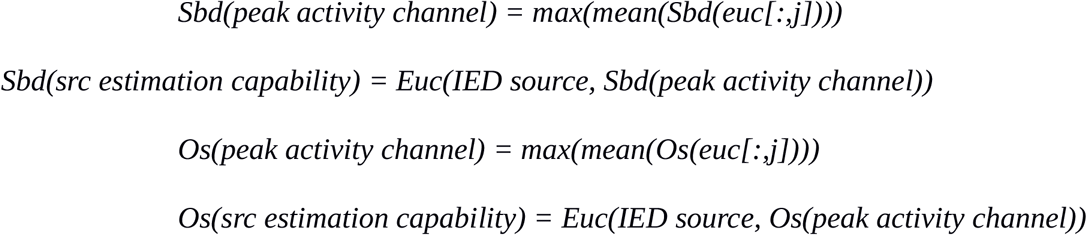
4. As a final step, a *dissimilarity score* comparing the source estimation capability of osMEG and sbdEEG was computed:

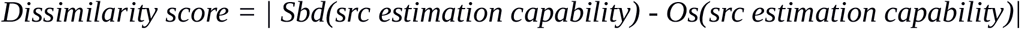

#### Source depth

The Euclidean distance between the center of the source and the adjacent electrode/sensor was quantified. Linear regression analysis was performed to determine whether source depth could predict dissimilarity score.

#### Source direction

Any influence of source direction on osMEG-sbdEEG similarity was analyzed in two ways.

1. The source direction was quantified as the angle between vector normal to the source, and the vector normal to the overlying gyral crown. Linear regression analysis was used to analyze any correlation between source direction and similarity score.
2. Secondly, sources that exhibited dissimilarity score 0 (low-DS sources) and sources that exhibited a dissimilarity score within the upper 15^th^ percentile (high-DS sources) were located and the orientation of the vector normal to the source center was determined. These orientations plotted as coordinates on the unit sphere. For all high-DS sources, all low-DS within the Euclidean distance 0.25 on the unit sphere were located. The distances between each high-DS source and the low-DS sources within this area was determined. A one-way ANOVA test was used to determine if any such region exhibited a statistically significant longer distance between high-DS and low-DS sources compared to all other regions. (see Fig 3 for an illustration of the source direction definitions)

**Figure 3:**
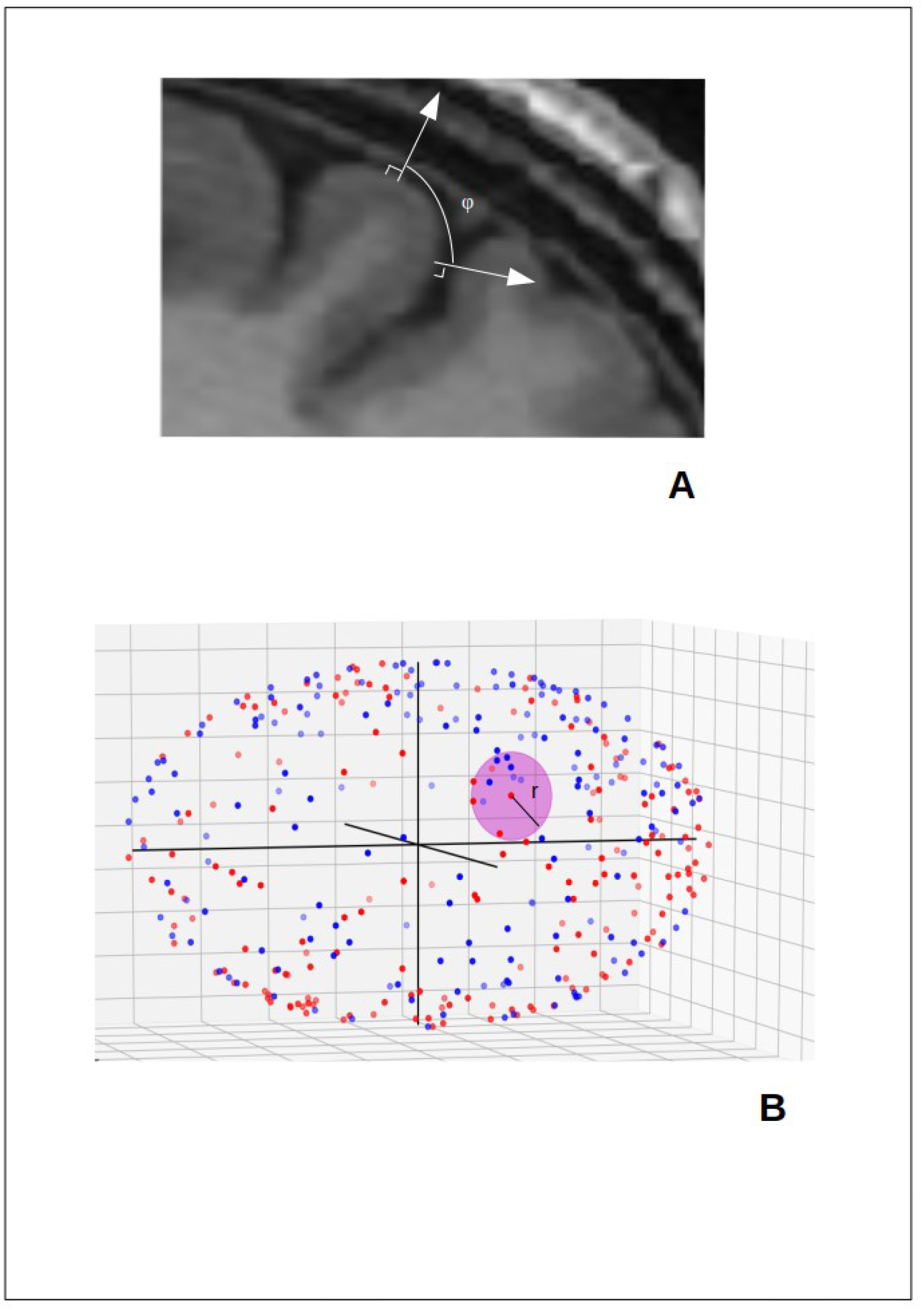
Definition of source directions. 3A: Source direction definition 1: Source directions defined as the angle phi between the vector normal to the interictal epileptiform discharge (IED) generating patch and the vector normal to the overlying gyral crown. 3B: Source direction definition 2: Low dissimilarity score (low-DS) (blue dots) and high-DS sources (red dots) with orientation of the normal vector plotted on a unit sphere. For all high-DS sources, all low-DS sources within an Euclidean distance of 0.25 (ua) were located (magnenta-colored patch, r = 0.25 (ua)).

## 3. Results

### 3.1 Irritative zone localization

Clinical IED source localization is based upon visual inspection of raw data and thereafter visually identifiable IEDs can be localized using ECD. Inspection of the raw sensor data of the gradually propagating epileptic foci demonstrated that the IEDs were visually identifiable first in the sbdEEG raw sensor data (mean focus size = 1 cm^2^) and secondly in the osMEG data (mean focus size = 2.2 cm^2^). Corresponding mean foci sizes for convMEG and hdEEG that gave rise to visually identifiable epileptic spikes were 20 cm^2^ and 21.6 cm^2^ respectively.

To further compare osMEG source estimation capability to other existing non-invasive techniques as well as to a sbdEEG whole-head grid, ECD of IED events were also performed regardless of raw sensor data visibility. As expected, sbdEEG exhibited superior source localization, where majority of dipoles were placed within the epileptic foci from mean foci size 0.8 cm^2^. For osMEG, the majority of dipoles were located within the epileptic foci from mean foci size 3.1 cm^2^. ConvMEG and hdEEG fitted dipoles were located within epileptic foci larger than 10 cm^2^ and 15 cm^2^ respectively. Consequently, osMEG dipoles were statistically significantly (p-value < 0.05) closer to source origin than hdEEG and convMEG dipoles for foci sizes between 3 and 23 cm^2^. SbdEEG dipoles were significantly closer to all sources except for lateral temporal and fronto polar sources compared to osMEG dipoles (p < 0.05). Below foci size 3 cm^2^, all four modalities performed at random level.

For details, see Table 1 and Figure 4.

**Table 1:**
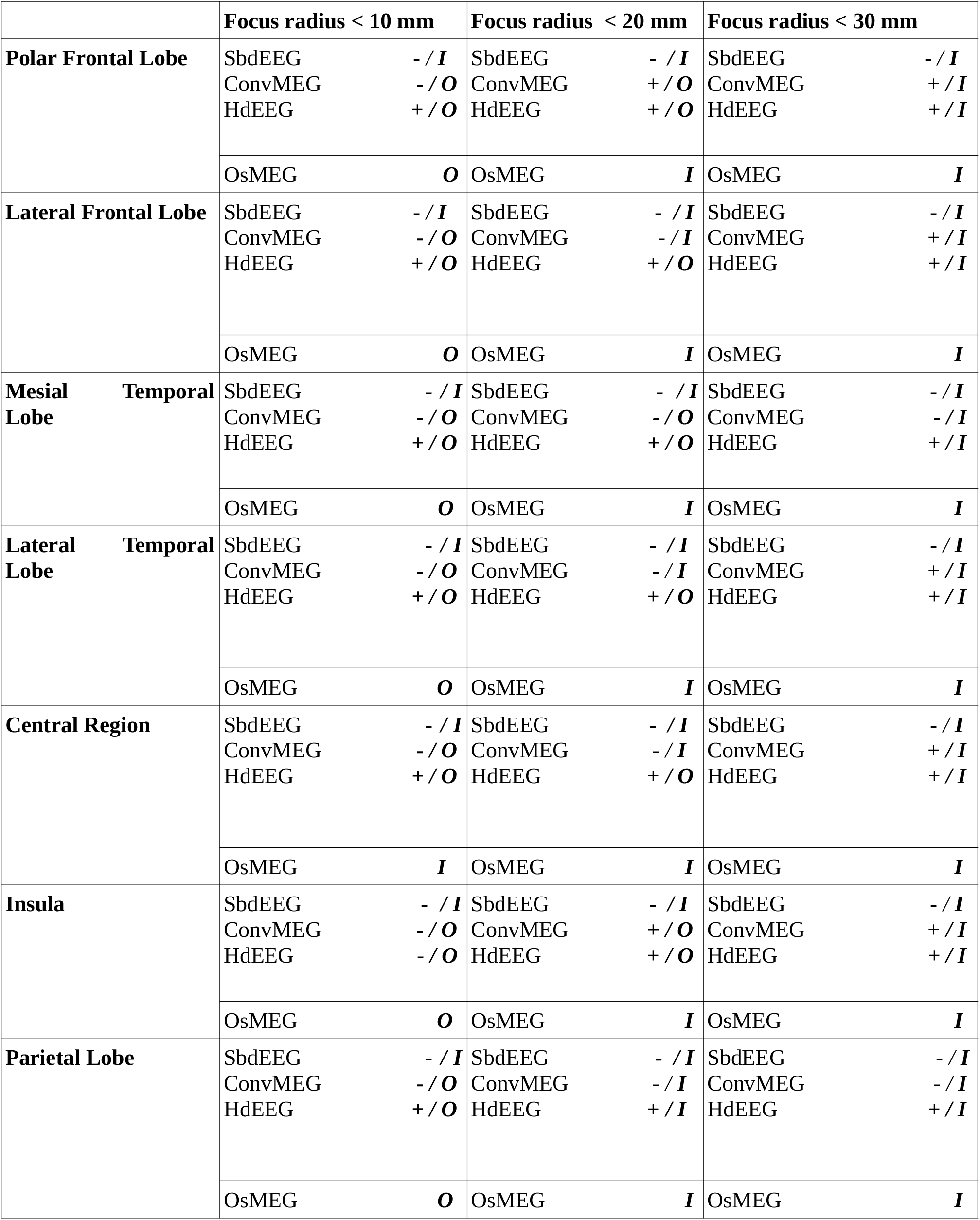
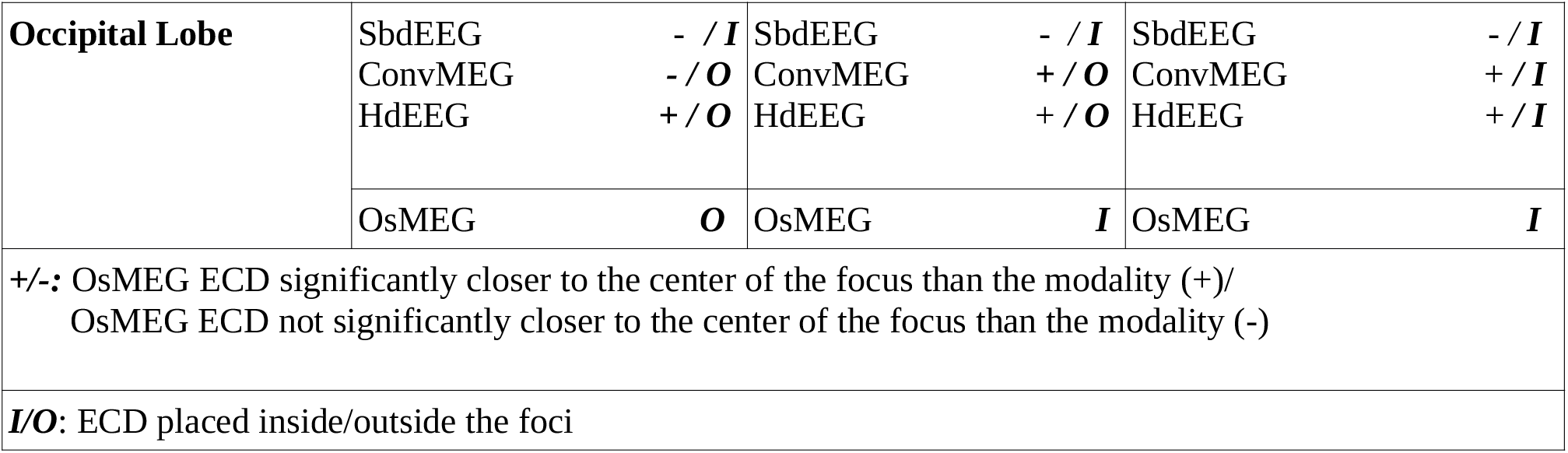
Comparison of the distance between center of growing epileptic focus and the equivalent current dipoles (ECDs) of subdural EEG (sbdEEG), conventional MEG (convMEG), high-density EEG (hdEEG) and on scalp MEG (osMEG) (+ : osMEG ECD significantly closer to the center of foci with the given radius; -: osMEG ECD not significantly closer to the center of foci with the given radius) and summary of whether the majority of ECDs of for the given foci sizes lie inside (***I*)** or outside (***O***) the focus.

|  | Focus radius < 10 mm |  | Focus radius < 20 mm |  | Focus radius < 30 mm |  |
| --- | --- | --- | --- | --- | --- | --- |
| **Polar Frontal Lobe** | SbdEEG | - / *I* | SbdEEG | - / *I* | SbdEEG | - / *I* |
|  | ConvMEG | - / *O* | ConvMEG | + / *O* | ConvMEG | + / *I* |
|  | HdEEG | + / *O* | HdEEG | + / *O* | HdEEG | + / *I* |
|  | OsMEG | *O* | OsMEG | *I* | OsMEG | *I* |
| **Lateral Frontal Lobe** | SbdEEG | - / *I* | SbdEEG | - / *I* | SbdEEG | - / *I* |
|  | ConvMEG | - / *O* | ConvMEG | - / *I* | ConvMEG | + / *I* |
|  | HdEEG | + / *O* | HdEEG | + / *O* | HdEEG | + / *I* |
|  | OsMEG | *O* | OsMEG | *I* | OsMEG | *I* |
| **Mesial Temporal Lobe** | SbdEEG | - / *I* | SbdEEG | - / *I* | SbdEEG | - / *I* |
|  | ConvMEG | - / *O* | ConvMEG | - / *O* | ConvMEG | - / *I* |
|  | HdEEG | + / *O* | HdEEG | + / *O* | HdEEG | + / *I* |
|  | OsMEG | *O* | OsMEG | *I* | OsMEG | *I* |
| **Lateral Temporal Lobe** | SbdEEG | - / *I* | SbdEEG | - / *I* | SbdEEG | - / *I* |
|  | ConvMEG | - / *O* | ConvMEG | - / *I* | ConvMEG | + / *I* |
|  | HdEEG | + / *O* | HdEEG | + / *O* | HdEEG | + / *I* |
|  | OsMEG | *O* | OsMEG | *I* | OsMEG | *I* |
| **Central Region** | SbdEEG | - / *I* | SbdEEG | - / *I* | SbdEEG | - / *I* |
|  | ConvMEG | - / *O* | ConvMEG | - / *I* | ConvMEG | + / *I* |
|  | HdEEG | + / *O* | HdEEG | + / *O* | HdEEG | + / *I* |
|  | OsMEG | *I* | OsMEG | *I* | OsMEG | *I* |
| **Insula** | SbdEEG | - / *I* | SbdEEG | - / *I* | SbdEEG | - / *I* |
|  | ConvMEG | - / *O* | ConvMEG | + / *O* | ConvMEG | + / *I* |
|  | HdEEG | - / *O* | HdEEG | + / *O* | HdEEG | + / *I* |
|  | OsMEG | *O* | OsMEG | *I* | OsMEG | *I* |
| **Parietal Lobe** | SbdEEG | - / *I* | SbdEEG | - / *I* | SbdEEG | - / *I* |
|  | ConvMEG | - / *O* | ConvMEG | - / *I* | ConvMEG | - / *I* |
|  | HdEEG | + / *O* | HdEEG | + / *I* | HdEEG | + / *I* |
|  | OsMEG | *O* | OsMEG | *I* | OsMEG | *I* |
| **Occipital Lobe** | SbdEEG | - / **I** | SbdEEG | - / **I** | SbdEEG | - / **I** |
|  | ConvMEG | - / **O** | ConvMEG | + / **O** | ConvMEG | + / **I** |
|  | HdEEG | + / **O** | HdEEG | + / **O** | HdEEG | + / **I** |
|  | OsMEG | **O** | OsMEG | **I** | OsMEG | **I** |
| +/-: OsMEG ECD significantly closer to the center of the focus than the modality (+)/<br>OsMEG ECD not significantly closer to the center of the focus than the modality (-) |  |  |  |  |  |  |
| **I/O**: ECD placed inside/outside the foci |  |  |  |  |  |  |

**Table 2.** The distance between the center of the seizure onset zone (SOZ) and the equivalent current dipoles (ECDs) of initial low-voltage, high-frequency seizure activity using subdural EEG (sbdEEG), on scalp MEG (osMEG), conventional MEG (convMEG) and high-density EEG (hdEEG).

|  | **SbdEEG ECD - center of SOZ, distance (mm)** | **OsMEG ECD - center of SOZ, distance (mm)** | **ConvMEG ECD - center of SOZ, distance (mm)** | **HdEEG ECD - center of SOZ, distance (mm)** |
| --- | --- | --- | --- | --- |
| **Mesial temporal lobe** | 2 | 12<br>(Correct lobe) | 52<br>(Incorrect lobe) | 45<br>(Incorrect lobe) |
| **Dorsal frontal lobe** | 3 | 4 | 5 | 8 |

**Figure 4:**
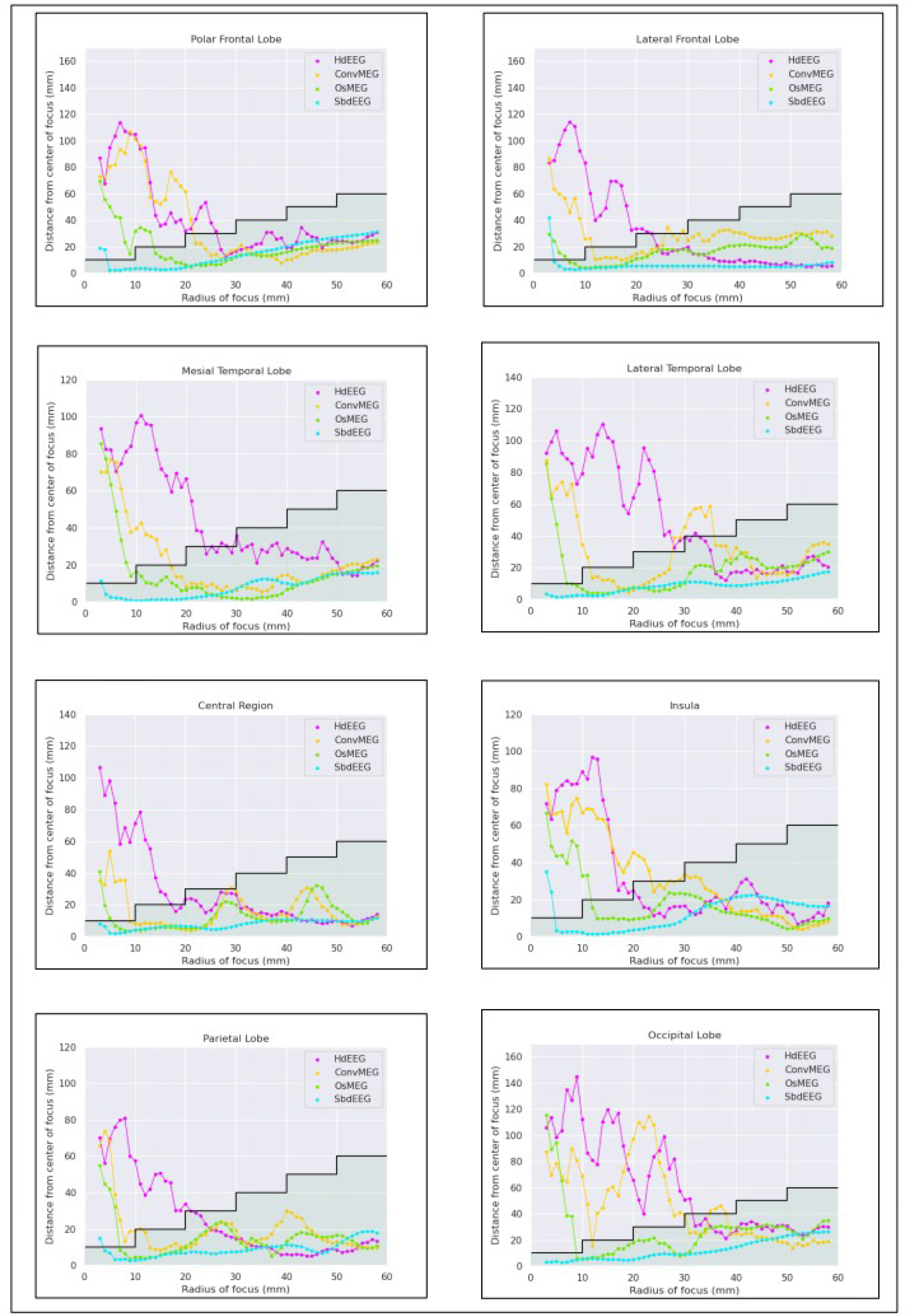
Equivalent current dipoles (ECDs) for all modalities across the eight selected brain regions. For each analyzed region, the distance between the center of the interictal epileptiform discharge (IED)-generating epileptic focus was plotted against the radius of the docus (range 3-60 mm). Green area below the step graph indicates ECDs placed within the focus. (SOZ: Seizure onset zone; HdEEG: High-density EEG, ConvMEG: conventional MEG, OsMEG: on scalp MEG, sbdEEG: Subdural EEG).

### 3.2 Seizure network localization

#### Mesial temporal lobe

SbdEEG localized the center of the SOZ with highest precision (9 mm SOZ center-dipole distance). OsMEG exhibited the second highest precision with 14 mm SOZ center-dipole distance. The osMEG dipole was placed within the mesial lobe. ConvMEG and hdEEG SOZ center-dipole distance were 70 mm and 80 mm, respectively. Both convMEG and hdEEG fitted dipoles were placed outside of the temporal lobe.

#### Dorsal frontal lobe

All modalities exhibited similar SOZ localization ability. The SOZ center-dipole distance was less than 1 cm for all modalities, and all modalities localized the SOZ to the correct sublobar region.

Please also see Figures 5a and 5b

**Figure 5a:**
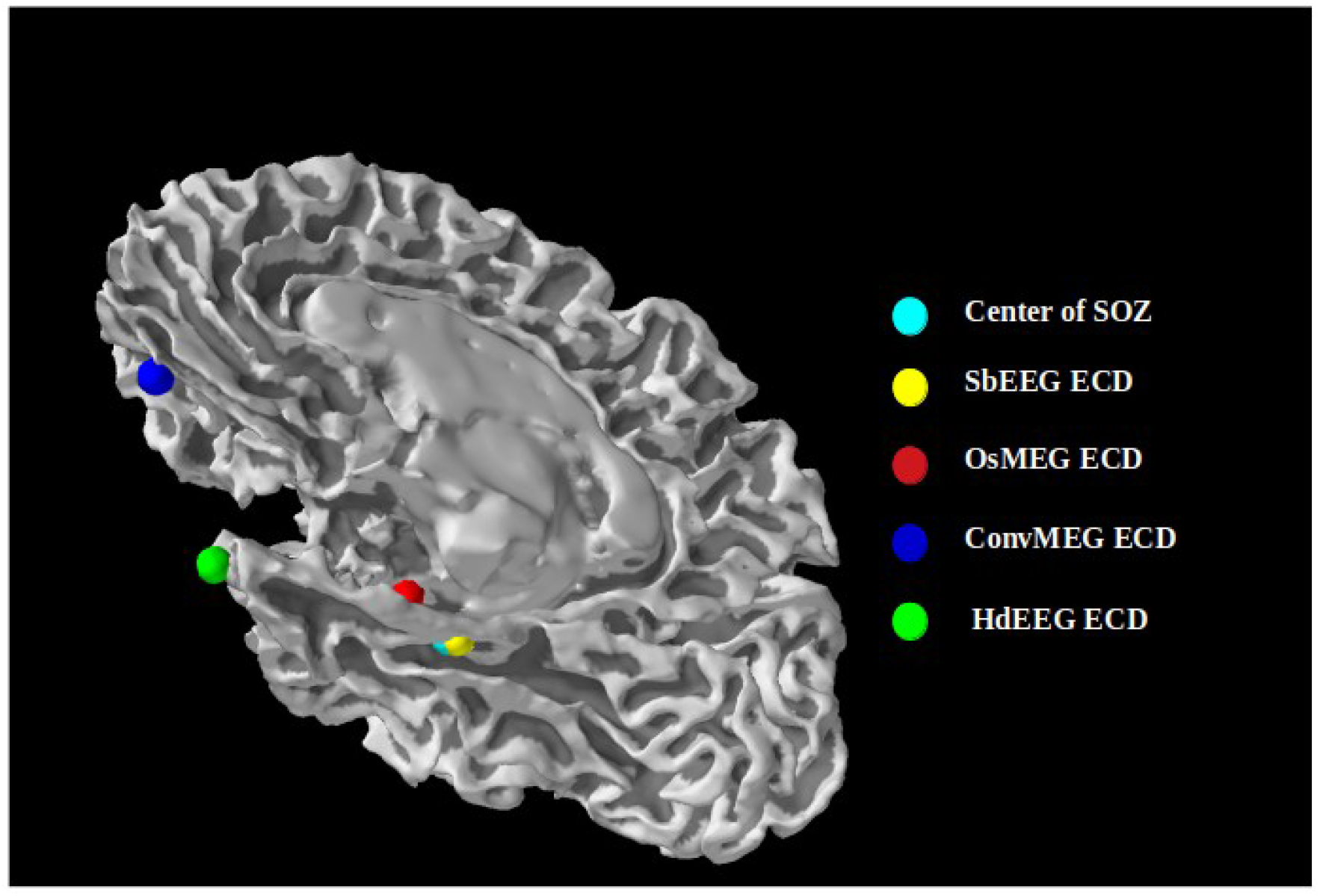
Equivalent current dipoles (ECDs) of mesial temporal lobe seizure onset zone (SOZ) for all modalities. Placement of the center of the seizure onset zone (SOZ) along side equivalent current dipoles (ECDs) of initial low-voltage, high-frequency seizure activity originating from mesial temporal lobe with rapid propagation to frontal lobe and contralateral temporal lobe determined using subdural EEG (sbdEEG), on scalp MEG (osMEG), conventional MEG (convMEG) and high-density EEG (hdEEG).

**Figure 5b:**
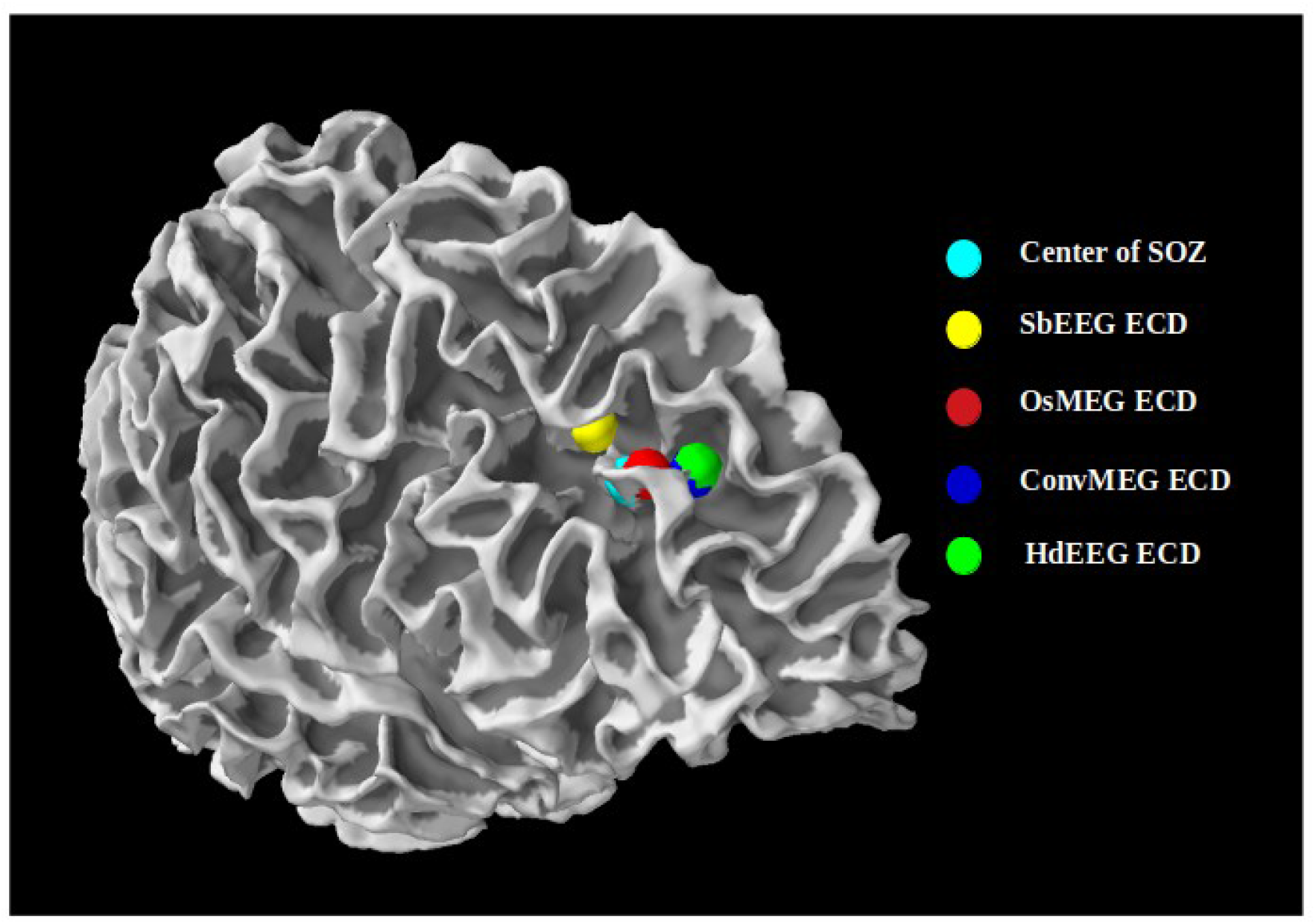
Equivalent current dipoles (ECDs) of lateral frontal lobe seizure onset zone (SOZ) for all modalities. Placement of the center of the seizure onset zone (SOZ) along side equivalent current dipoles (ECDs) of initial low-voltage, high-frequency seizure activity originating from lateral frontal lobe with propagation to contralateral frontal lobe determined using subdural EEG (sbdEEG), on scalp MEG (osMEG), conventional MEG (convMEG) and high-density EEG (hdEEG).

### 3.3 Cancellation index

There was no statistically significant difference between osMEG cancellation indices for the eight epileptic foci. There were neither any statistically significant difference between osMEG and convMEG cancellation indices for any epileptic foci. The cancellation index of both convMEG and OsMEG were significantly higher than that of hdEEG for all epileptic foci (p-value < 0.5)

### 3.4 Comparison of sbdEEG and osMEG epilepsy activity detection

Comparing how much sbdEEG and osMEG source localization differs for all locations across the right hemisphere, 12.78% exhibited identical sbdEEG-osMEG source localization. Median dissimilarity score was 2.2 mm, and the maximum dissimilarity score was 44 mm (see Fig 6). Analyzing the importance of source depth demonstrate a weak dependence of source depth (similarity score: linear regression coefficient = .034 and intercept = .002). However, linear regression analysis demonstrated no dependence on source orientation (linear regression coefficient = −1.3e-06 and intercept = .004). No specific source orientation exhibited a statistically significant longer distance between high-DS and low-DS sources compared to all other source orientations.

**Figure 6:**
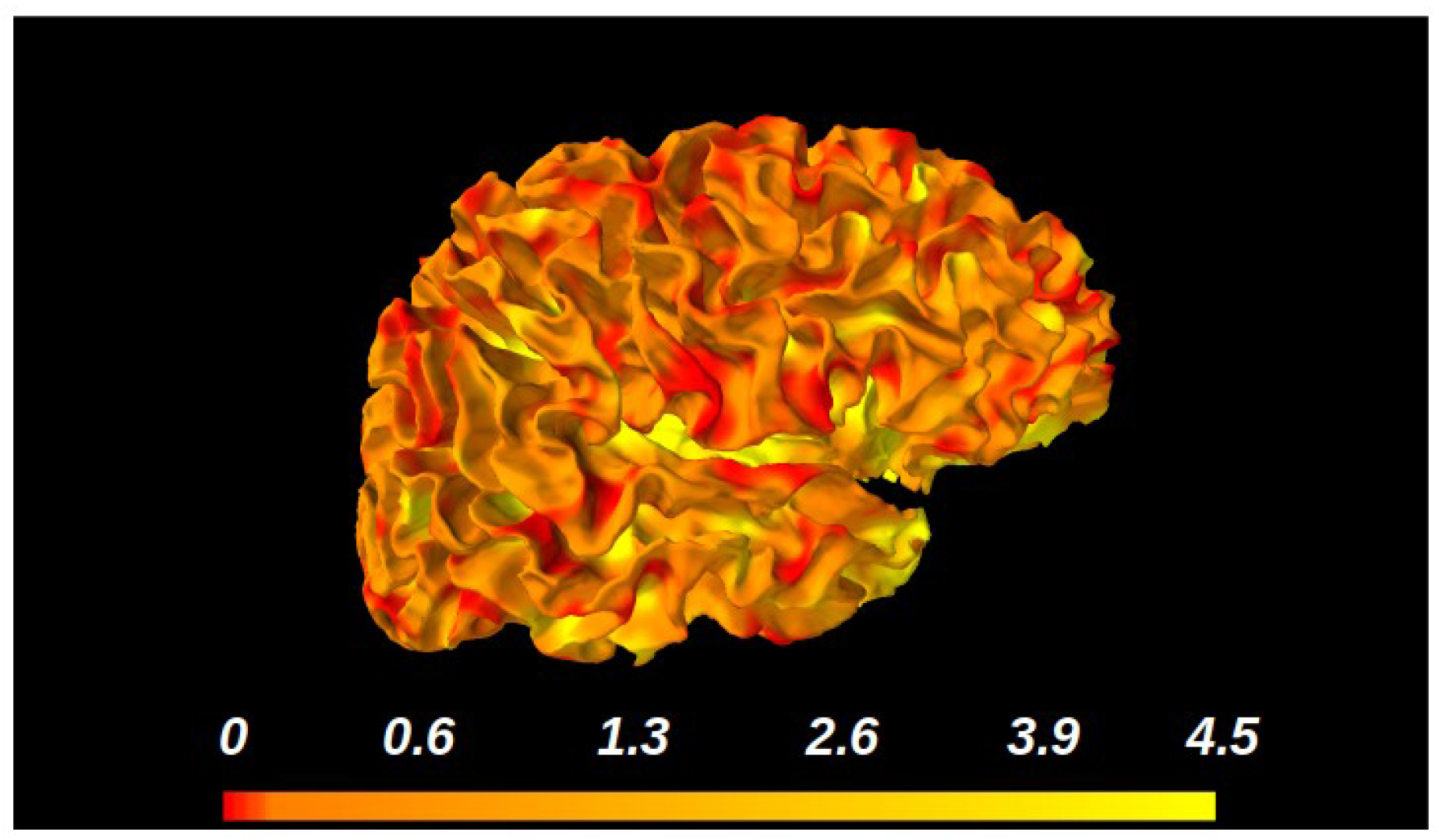
On scalp MEG (osMEG) and subdural EEG (sbdEEG) dissimilarity scores. Difference (unit: centimeter) between interictal epileptiform discharge (IED) source estimation of osMEG and sbdEEG (for definition of the dissimilarity score, see section *2.6 Comparison of osMEG and sbdEEG*) across the cortical mantle (radius of IED-generating cortical patch = 5 mm).

## 4. Discussion

This study aimed to characterize osMEG performance in a simulated realistic epilepsy evaluation, by examining the capacity of osMEG to detect and localize simulated seizures and interictal epileptic spikes, and by comparing osMEG capabilities to those of convMEG, hdEEG and sbdEEG. To our knowledge, this is the first-ever osMEG modelling study to also include an hdEEG comparison.

The results show that osMEG is well suited to detect and estimate the source of epileptic activity. OsMEG is superior to both convMEG and hdEEG, and succeeds in localization of small seizure onset zones with an accuracy comparable to that of sbdEEG. OsMEG detected IEDs as well as seizures from smaller epileptic foci than did both convMEG and hdEEG.

The increased signal strength of osMEG as compared to convMEG and hdEEG enabled improved source localization of both low amplitude seizure onset zone activity, and interictal epileptic spikes. Importantly, we demonstrated that the whole-head osMEG sensor system can correctly localize rapidly propagating mesial temporal lobe seizures that are difficult to register with hdEEG and that are difficult to estimate using both convMEG and hdEEG. Thus, our results indicate that osMEG is able to detect and localize seizures that would otherwise require sbdEEG, however non-invasively, and with whole-head coverage. These findings are in accordance with previous osMEG sensor studies.

To support these modelling results, we recently performed the first ever osMEG measurement on an epilepsy patient and demonstrated that osMEG was capable of detecting twice as many interictal epileptic spikes than convMEG, half of which were low-amplitude interictal epileptic spikes not seen by either convMEG nor EEG (Westin et al., 2020). In further agreement with our results, experimental osMEG studies have demonstrated that osMEG sensors generate a higher signal amplitude in general as compared to convMEG, and that osMEG spatial resolution is superior to that of convMEG (Andersen et al.; Xie et al., 2015; Boto et al., 2021).

From the perspective of clinical utilization, we here demonstrated that osMEG epileptic activity localization ability dramatically exceeds that of hdEEG, indicating that osMEG can provide improved non-invasive neurophysiological epilepsy evaluations. In this simulation study, we had whole-brain coverage with sbdEEG, which is impossible in reality, as the whole brain cannot be implanted, and the yield of sbdEEG largely depends on the accuracy of the hypothesis the impantation is based on. While sbdEEG covers only smaller subregions, the osMEG whole head coverage enables measurement of co-existing, separately located seizure generating zones.

As our results indicate that osMEG epileptic activity detection and localization ability was superior to that of convMEG and hdEEG and second only to sbdEEG, we also evaluated sbdEEG-osMEG source localization dissimilarity. These results showed that the mean distance between sbdEEG and osMEG source localization was less than 10 mm, which shows that the osMEG spatial resolution resembles intracranial registrations. However, the comparison also demonstrated that osMEG was less sensitive to deep sources than was sbdEEG. This is in accordance with previous studies indicating that osMEG spatial resolution is depth dependent (Iivanainen et al., 2017). Interestingly, we did not find any difference in source orientation sensitivity between osMEG and sbdEEG, indicating that osMEG might be able to detect activity from source orientations not seen by convMEG. This notion is in agreement with experimental osMEG data from our group, showing that osMEG detects radial activity undetectable by convMEG. Since osMEG sensors are positioned directly on scalp, the angles between the cortical sources and sensors change compared to convMEG. Thus, it is possible that a whole-head osMEG sensor system give rise to fewer true radial sources than do convMEG.

### Limitation

There are several limitations to this study. Most importantly, a whole-head osMEG recording on epileptic patients has yet to be performed. Thus, there are no real measurements or quantifications of osMEG epileptic brain noise amplitudes. Especially, epilepsy patients often exhibit generalized or focal pathological slowing, sometimes of high amplitude. It is unknown how such activity would be detected by osMEG sensors. Thus, any estimation of irritative zone brain noise would be speculative. As a simplification, background brain noise levels were set equal to convMEG. Additional research and including experimental studies is warranted in order to quantify realistic epilepsy brain noise levels. Furthermore, ECD was used to compare localization of extended sources. ECD is mathematically unsuited for larger sources. There exist several inverse solutions dedicated to extended sources but none of these have been clinically validated why ECD (which has been clinically validated) was chosen nonetheless (Jerbi et al., 2004).

### Conclusion

This study indicate that a whole-head osMEG sensor system would be capable of very accurate non-invasive source localization, better than both hdEEG and convMEG, and close to the performance of sbdEEG (while also having the benefits of being non-invasive and allowing whole-head coverage). OsMEG thus appears to have a very promising potential to become a safe, high-resolution technique for advanced non-invasive, whole head epilepsy evaluations.

## Abbrevations

osMEG: on scalp magnetoencephalography
convMEG: conventional magnetoencephalography
hdEEG: high density electroencephalography
sbdEEG: subdural electroencephalography
IED: interictal epileptiform discharge
OPM: optically pumped magnetometer
highTc SQUID: high critical temperature superconducting quantum interference devices
SOZ: seizure onset zone

